# The epigenetics effects of transposable elements are context dependent and not restricted to gene silencing

**DOI:** 10.1101/2023.11.27.568862

**Authors:** Marta Coronado-Zamora, Josefa González

## Abstract

Transposable elements (TEs) represent a threat to genome integrity due to their proliferation capacity. Eukaryotic cells silence TEs through different epigenetic mechanisms, including the deposition of repressive histone marks. Previous studies have shown that repressive marks can spread to neighboring sequences. However, evidence for this spreading affecting nearby gene expression remains limited. Similarly, whether TEs induce changes in the enrichment of active histone marks genome-wide, and its potential impact on gene expression have not been widely studied. In this work, we performed a comprehensive study of the epigenetic effects of 2,235 TEs and their potential effects on nearby gene expression on *D. melanogaster* head, gut and ovary. While most of the TEs (816) induce the enrichment of the H3K9me3 repressive mark, with stronger epigenetic effects in the ovary, a substantial number (345 TEs) induce the enrichment of the H3K27ac active mark, particularly in the gut. We found that 70% of the H3K9me3 enriched TEs induced gene down-regulation, and 50% of the H3K27ac enriched TEs induced gene up-regulation. These changes in expression affect specific regulatory networks in head and gut while in ovary, genes were not enriched for any biological functions. Furthermore, TE epigenetic effects on gene expression are genomic context dependent. Finally, we found that TEs also affect gene expression by disrupting regions enriched for histone marks. Overall, our results show that TEs do generate regulatory novelty through epigenetic changes, with these epigenetic effects not restricted to gene silencing and being context dependent.

**Significance statement:** Transposable elements (TEs) are repetitive DNA sequences found in nearly all studied organisms that have the capacity to move within the genome. To prevent their proliferation, eukaryotic cells target TEs with repressive histone marks, an epigenetic signal that blocks their expression. While these repressive marks can spread to neighboring genes, the evidence of how this impacts gene expression is limited. Similarly, whether TEs also influence the enrichment and depletion of active histone marks and their genome-wide impact is not understood. In this work, we studied the histone mark enrichment of 2,235 polymorphic TEs across three body parts of *D. melanogaster*. Our results provide evidence for the genome-wide role of TEs in the generation of regulatory novelty through epigenetic changes.

## INTRODUCTION

The capacity of transposable elements (TEs) to move and expand within the genome is a threat to genome integrity (1, 2). As a result, cells have evolved a range of molecular strategies, including epigenetic mechanisms, to repress and control TE activity (3). One epigenetic mechanism widespread among eukaryotes is the methylation of H3K9 (H3K9me3), a repressive mark associated with the formation of heterochromatin, that targets TEs to suppress their activity (4–7). While the enrichment of repressive epigenetic marks at TEs would limit the expression of TEs, which should be beneficial for the cell, several studies have shown that TE silencing could also influence nearby gene sequences across species, including plants (8–10) and animals (5, 11, 12). A potential functional consequence of the spread of repressive marks is the down-regulation of the expression of nearby genes. However, the evidence for this effect genome-wide is limited (1, 13, 14). The few studies comparing the expression of alleles with and without TE insertions with epigenetic effects found no clear association between enrichment of H3K9me3 and gene down-regulation (13, 15, 16). Thus, whether the spreading of H3K9me3 from TEs to nearby gene regions results in gene down-regulation genome-wide is still an open question.

While TEs enrichment in H3K9me3 has been widely studied across taxa, less is known about the enrichment of active histone marks in TE sequences. Some studies, mainly using human data, have found evidence of TEs inserted in active chromatin-state regions suggesting that these TEs could be acting as *cis*-regulatory elements (17–20). However, the majority of analyses available are based on single-genomes, and could not discern whether TEs induced the enrichment of active marks or were inserted into regions that were already enriched for those marks. Indeed, the few studies comparing alleles with and without TE insertions found that at least some TE families preferentially insert into genomic regions enriched for histone marks (21–23). In *Drosophila,* there is only evidence for an individual TE insertion that, under oxidative stress conditions, recruits an active epigenetic mark, inducing changes in the expression of adjacent genes (24). Therefore, our understanding of how TEs contribute to the genome-wide enrichment of active marks is still very limited. Similarly, whether TE insertions disrupt the epigenetic landscape of the regions where they insert has not been widely explored. There is evidence in *D. melanogaster* for the depletion of active marks in the regions flanking TE insertions (15). However, whether this pattern is TE-induced, and whether it impacts gene expression has not been investigated.

In this work, we performed a thorough study of the epigenetic effects of 2,235 TEs and their potential effects on nearby gene expression using ChIP-seq data, for a repressive and an active histone mark, and transcriptomic data obtained from head, gut and ovary of five *D. melanogaster* strains. These five strains have high quality *de novo* reference genomes and TE annotations available, which allowed us to systematically characterize the epigenetic effect of TEs (25, 26). We identified TE insertions that induce epigenetic changes, both enrichment and depletion of the repressive H3K9me3 and the active H3K27ac histone marks, study whether the epigenetic effect of these TEs was body part-specific, assess whether these effects were induced by particular TE families, and evaluate the potential functional implications of TE-induced epigenetic effects on gene expression.

## RESULTS

### 1. TEs induce the enrichment of active and repressive histone marks differentially across body parts

To gain insight into the epigenetic effects of TEs across adult body parts, we took advantage of the high quality *de novo* TE annotations and ChIP-seq data, produced for the repressive mark H3K9me3 and for the active mark H3K27ac, in head, gut and ovary, from five *D. melanogaster* European strains (25, 26). We first analyzed the average epigenetic states of 4,823 TE flanking sequences (±20kb) across the three body parts. In the adult head, we observed an average depletion of the repressive mark H3K9me3, which contrasts with the findings of Lee and Karpen (2017) (15), who reported a weak enrichment in the head (**Fig. 1A** and **Table S1A**). In the gut and ovary, there is a strong average enrichment of H3K9me3 in the TE flanking sequences, as previously described (15) (**Fig. 1A**). Regarding the active mark, all three body parts showed a depletion of H3K27ac, supporting previous observations that active histone modifications are depleted near TE insertions genome-wide (15) (**Fig. 1A** and **Table S1A**). However, by analyzing the average epigenetic effects of all TEs annotated in the genomes studied, we cannot discard that these patterns are due to the preferential insertion of TEs in regions enriched or depleted for these histone marks (21–23, 27).

**Figure 1.**
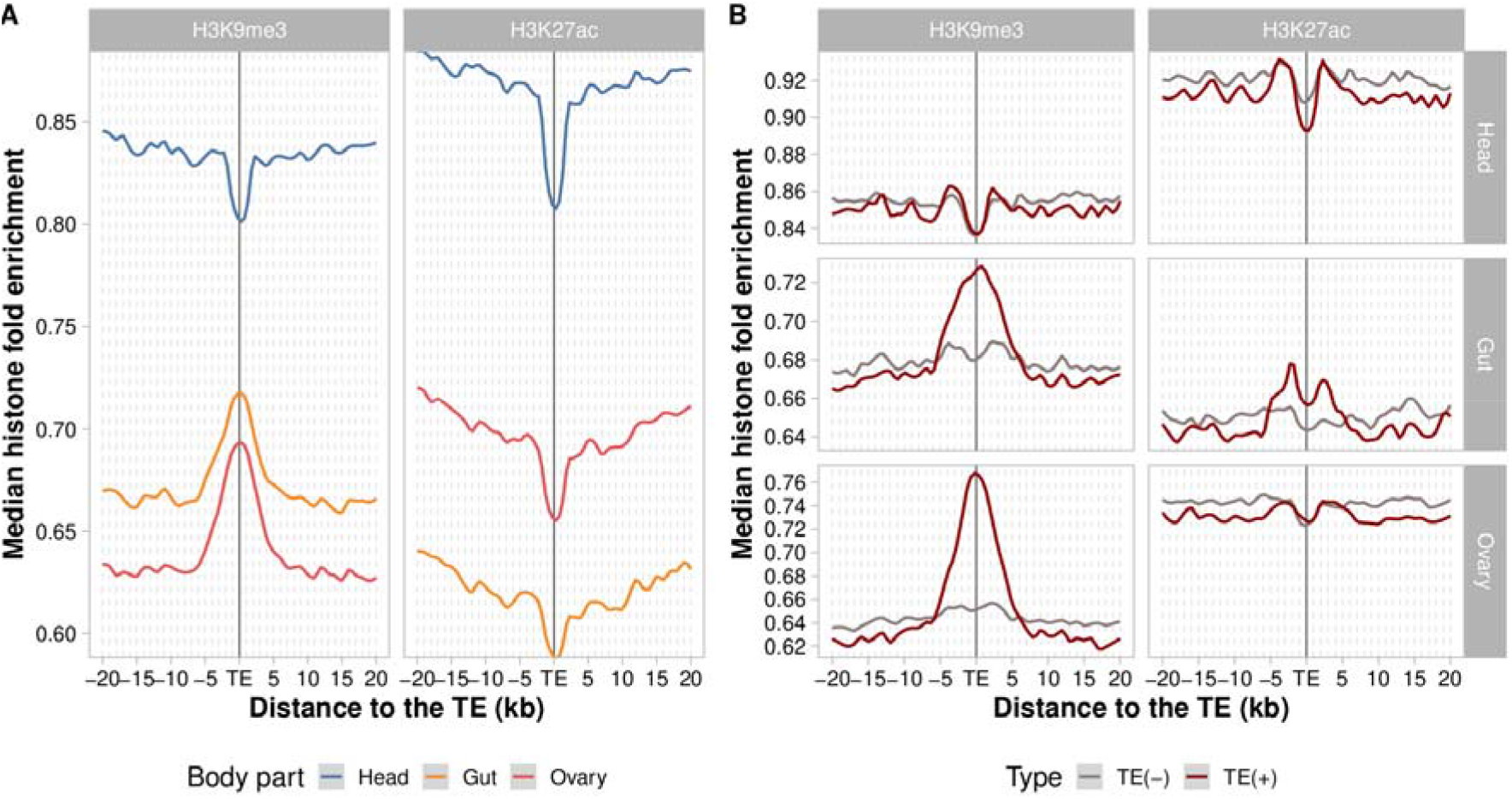
Epigenetic states in the ±20kb TEs flanking region across body parts. A. Median H3K9me3 and H3K27ac fold enrichment of all TEs annotated in five genomes (*n*=4,823 TEs). Head (blue line), gut (orange line) ovary (red line). B. Median H3K9me3 and H2K27ac fold enrichment of polymorphic TEs (*n*=2,325 TEs). Red line represents histone mark enrichment of genomes with a TE [TE(+)] and the gray line represents histone mark enrichment of genomes without a TE [TE(-)]. H3K9me3 and H3K27ac fold enrichment was averaged (median) over all sequences flanking the analyzed TEs of all genomes. Plots were generated using LOESS smoothing (span = 10%).

To test whether the observed enrichment and depletion of histone marks is TE-induced, we focused on polymorphic TEs, for which we can compare the epigenetic states of the TE flanking regions in genomes with and without the TE insertions. We performed this analysis for 2,325 polymorphic TEs (see *Materials and Methods*).

In the head, we did not observe differences between the genomes with and without TEs, suggesting that TEs are not driving the previously observed depletion in the H3K9me3 mark (**Fig. 1B** and **Table S1B**). On the other hand, TE insertions were associated with H3K9me3 enrichment in gut and ovary (6.4% and 16.6% enrichment in the flanking ±1kb window, respectively) (**Fig. 1B**). The lower levels of H3K9me3 observed in the head could be explained by the level of expression of the methyltransferases and demethylases. We found that the expression level of *Su(var)3-9*, a key heterochromatin regulator of H3K9me3, was lower in the head compared to gut and ovary (Wilcoxon test, *p*-values < 0.001 for both comparison, **Fig. S1**). Indeed, the expression level of Su(var) proteins has been already associated with the magnitude of TE’s epigenetic effects in *Drosophila* (Lee and Karpen 2017). On the other hand, the expression level of the *Kdm3* and *Kdm4B* demethylases were higher in the head compared to gut and ovary (Wilcoxon test, p-values < 0.05 for all comparisons, for *Kdm4B,* the comparison was only significant for head *vs* gut, **Fig. S1**). Regarding H3K27ac, we observed a slight depletion in genomes with TEs in head (−1.4%), an enrichment in gut (2.3%), and no enrichment in ovary (**Fig. 1B** and **Table S1B**). This contrasts with the previous observation of a general depletion of the active mark across body parts (**Fig. 1A**). Note that the enrichment pattern observed in gut is the one expected for active promoters, where the histone mark enrichment flanks rather than directly overlap with the enhancer sequences (28) (**Fig. 1B**).

Although the enrichment/depletion patterns described above (Fig. 1B) are only found in the genomes that contain the TE insertion, to discard that these patterns could be due to strain-specific epigenetic variation, we checked whether similar patterns were found in genomic regions that do not contain TE insertions (see Material and Methods). We did not find consistent patterns of enrichment or depletion in our TE-free genomic regions (Fig. S2) suggesting that the previously observed patterns (Fig. 1B) are most likely TE-induced.

### 2. Individual TEs mostly enrich for repressive marks in the ovary and for active marks in the gut and deplete similarly across histones and body parts

While overall, polymorphic TEs induce the enrichment and depletion of repressive and active marks across body parts (**Fig. 1B**), the analysis of individual TE insertions allows us to gain further insight into both the percentage of enrichment and depletion and the spread of these effects to nearby regions. We considered an individual TE to have an epigenetic effect if the fold enrichment of H3K9me3, H3K27ac, or both marks (bivalent) was significantly higher or lower within ±1kb of the TE insertion (see *Material and Methods*). For these TEs, we also computed: *i)* the percentage change in enrichment or depletion compared to strains without the TE within the ±1kb window, and *ii)*, the extent of spread from the TE insertion (up to ±20kb) with consecutive significant changes in histone marks. We computed the same metrics for our dataset of TE-free regions to show that these effects are induced by TEs and not because of epigenetic variability present in the strains (Table S2A).

We found that most of the polymorphic TEs analyzed exert epigenetic effects in at least one of the body parts analyzed (71.5%, 1,597/2,235; **Supplemental Data** and **Table S2B-C**). Among the TEs with epigenetic effects, 65.4% (1,045/1,597) induced the enrichment of one or both histone marks simultaneously (bivalent), 25% (400/1,597) induced the depletion of one or both histone marks, and 9.6% (152/1,597) induced the enrichment of histone marks in some body parts but the depletion in others. If we focus on the TEs that induce the enrichment of histone marks in at least one body part, we found that the most prevalent and strongest effect is the enrichment of H3K9me3: 816 out of 1,197 TEs (**Table 1** and **Fig. 2A**). The majority of these TEs exert their effect on the ovary (576/816), where the average percentage of enrichment and the spread of the histone mark is higher than in other body parts: 119.83% average enrichment (±100.39) and 3.76kb spread (±3.5) (Wilcoxon’s test, *p*-values < 0.001, **Fig. 2B-C**, **Table 1** and **Table S2D**). Head, in contrast, is the body part with less TEs inducing the enrichment of H3K9me3 compared to gut and ovary (140 TEs, χ² test, *p*-values < 0.001). Although we observed an average enrichment of H3K27ac only in the gut (**Fig. 1B**), we found that TEs also enrich for the active mark in other body parts (**Fig. 2A**). From 345 TEs that induce the enrichment of the active mark, half of them exert their effect in the gut (183/345), which is a percentage significantly higher than the TEs found in the ovary (112 TEs, χ² test, *p*-value < 0.001) and the head (88 TEs χ² test, *p*-value < 0.001) (**Fig. 2A**). The percentage of enrichment and spread is higher in the gut: 37.16% (±21.97) and 2.92kb (±2.95), respectively (Wilcoxon’s test, *p*-values < 0.05, **Fig. 2B-C**, **Table 1** and **Table S2D**). We found that the 183 TE insertions that enrich for H3K27ac in the gut were enriched for full-length copies (90 full-length *vs.* 93 fragment) compared to the TE insertions not enriched for H3K27ac in the gut (719 TEs, 264 full-length and 457 fragments; Fisher’s exact test, *p*-value < 0.01), suggesting that some of these insertions might be active. We also identified 304 TEs that induced the enrichment of both marks simultaneously (bivalent) (**Fig. 2A**). This effect is observed in similar proportions in gut (147 TEs) and ovary (133 TEs) (χ² test, *p*-value = 0.29) with stronger effects in the ovary than in the gut (Wilcoxon’s test, *p*-values < 0.05; **Fig. 2B-C**, **Table 1** and **Table S2D**).

**Figure 2.**
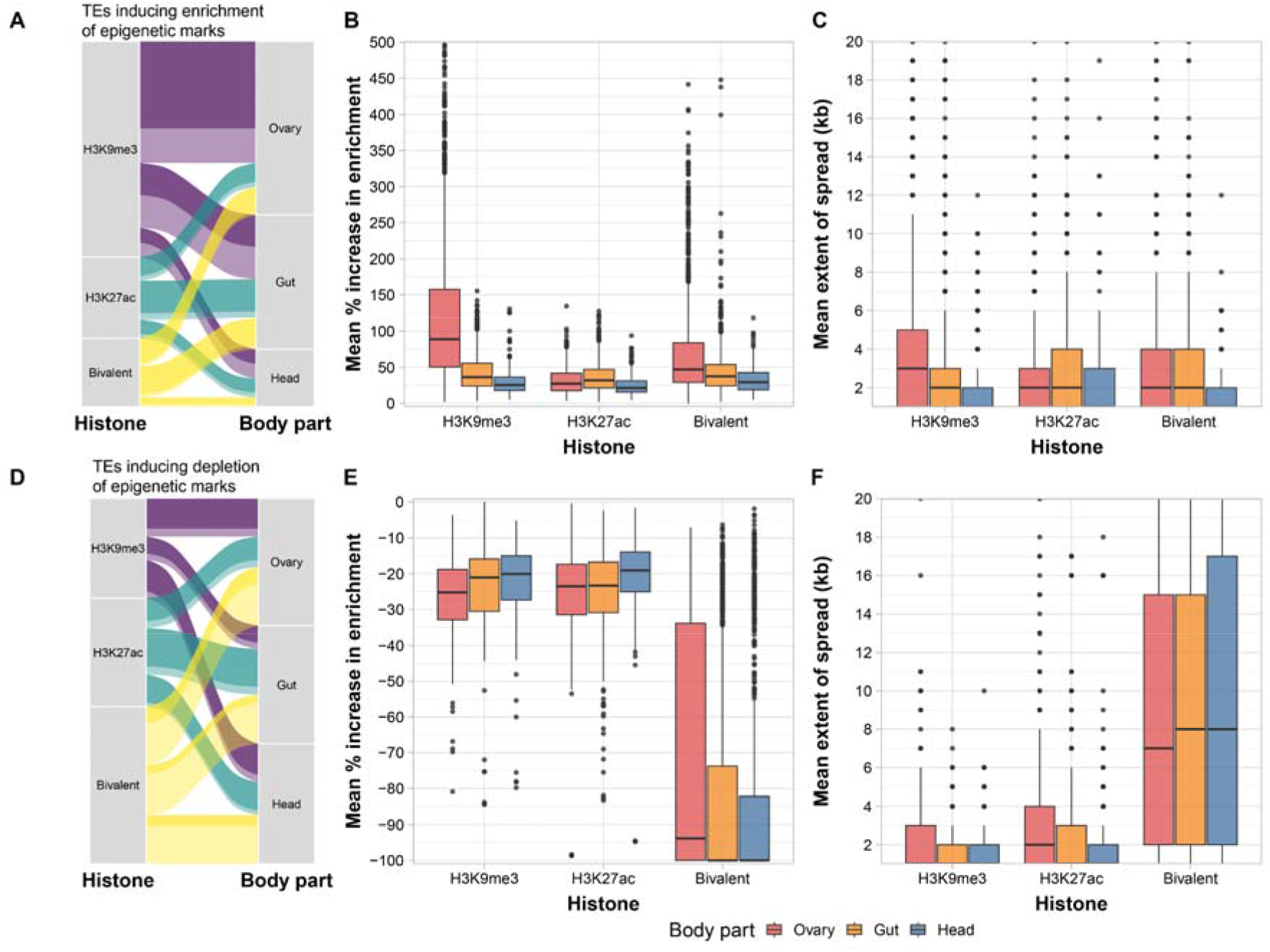
Epigenetics effects of 1,597 TEs across body parts. A. 1,197 TEs induce the enrichment of epigenetic marks. Dark colors indicate body part-specific TEs and light colors indicate TEs that are not body part-specific. **B. Mean percentage of increase in H3K9me3, H3K27ac and bivalent enrichment of the TEs across body parts.** Outliers removed (TEs with epigenetic effects > 500%) for clarity. **C. Mean extent of spread (kb) of H3K9me3, H3K27ac and bivalent enrichment) by body parts.** The spread was calculated in windows of up to 20kb adjacent to the TE insertion. **D. 552 TEs inducing the depletion of epigenetic marks.** Dark colors indicate body part-specific TEs and light colors indicate TEs that are not body part-specific. **E. Mean percentage of increase in H3K9me3, H3K27ac and bivalent enrichment of the TEs by body parts. F. Mean extent of spread (kb) of H3K9me3, H3K27ac and bivalent depletion of the TEs by body parts.** Values and statistics in Table 1 and Table S2.

**Table 1.**
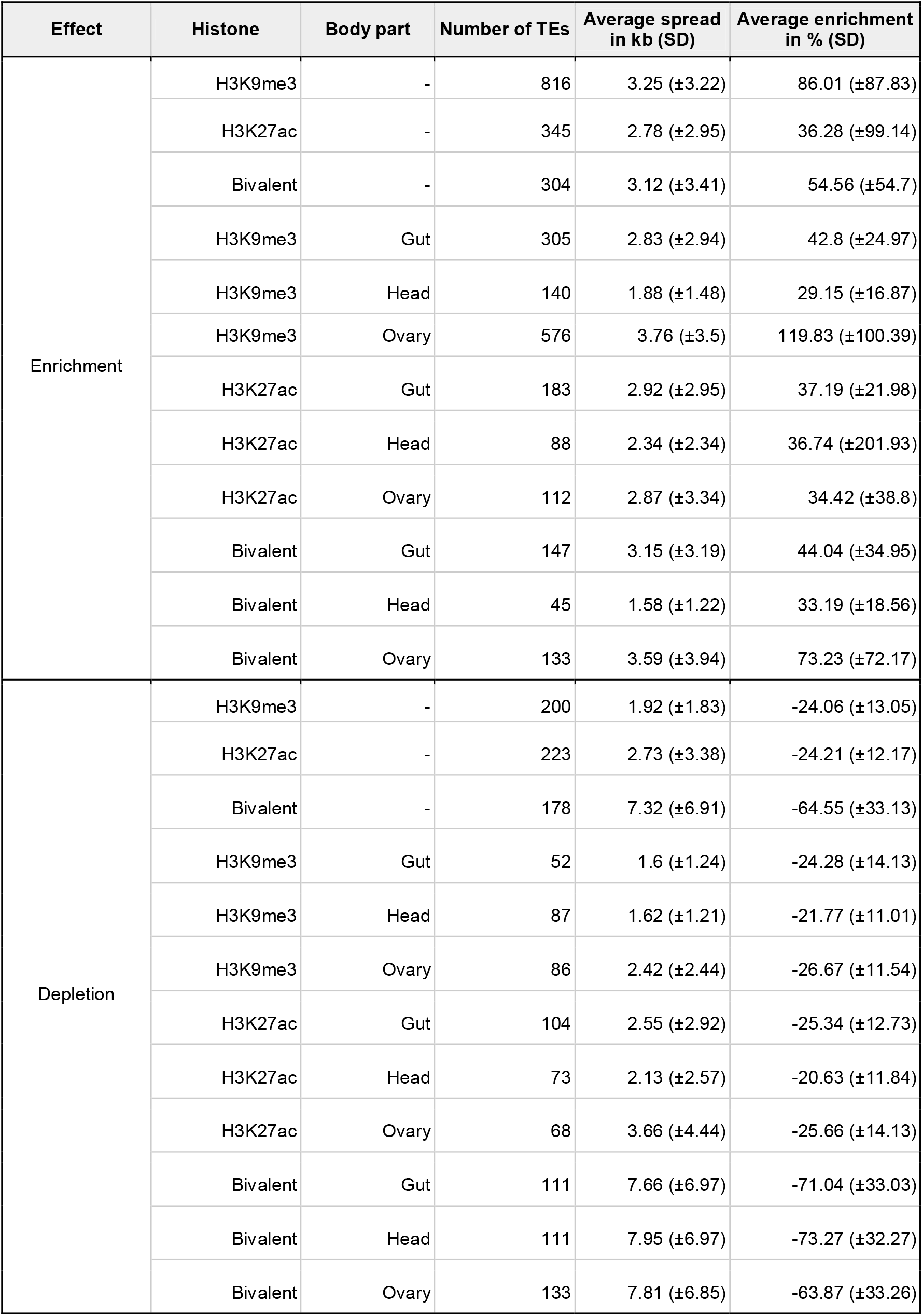
Epigenetic effects of 1,597 TEs.

| Effect | Histone | Body part | Number of TEs | Average spread in kb (SD) | Average enrichment in % (SD) |
| --- | --- | --- | --- | --- | --- |
| Enrichment | H3K9me3 | - | 816 | 3.25 ($\pm 3.22$) | 86.01 ($\pm 87.83$) |
| | H3K27ac | - | 345 | 2.78 ($\pm 2.95$) | 36.28 ($\pm 99.14$) |
| | Bivalent | - | 304 | 3.12 ($\pm 3.41$) | 54.56 ($\pm 54.7$) |
| | H3K9me3 | Gut | 305 | 2.83 ($\pm 2.94$) | 42.8 ($\pm 24.97$) |
| | H3K9me3 | Head | 140 | 1.88 ($\pm 1.48$) | 29.15 ($\pm 16.87$) |
| | H3K9me3 | Ovary | 576 | 3.76 ($\pm 3.5$) | 119.83 ($\pm 100.39$) |
| | H3K27ac | Gut | 183 | 2.92 ($\pm 2.95$) | 37.19 ($\pm 21.98$) |
| | H3K27ac | Head | 88 | 2.34 ($\pm 2.34$) | 36.74 ($\pm 201.93$) |
| | H3K27ac | Ovary | 112 | 2.87 ($\pm 3.34$) | 34.42 ($\pm 38.8$) |
| | Bivalent | Gut | 147 | 3.15 ($\pm 3.19$) | 44.04 ($\pm 34.95$) |
| | Bivalent | Head | 45 | 1.58 ($\pm 1.22$) | 33.19 ($\pm 18.56$) |
| | Bivalent | Ovary | 133 | 3.59 ($\pm 3.94$) | 73.23 ($\pm 72.17$) |
| Depletion | H3K9me3 | - | 200 | 1.92 ($\pm 1.83$) | -24.06 ($\pm 13.05$) |
| | H3K27ac | - | 223 | 2.73 ($\pm 3.38$) | -24.21 ($\pm 12.17$) |
| | Bivalent | - | 178 | 7.32 ($\pm 6.91$) | -64.55 ($\pm 33.13$) |
| | H3K9me3 | Gut | 52 | 1.6 ($\pm 1.24$) | -24.28 ($\pm 14.13$) |
| | H3K9me3 | Head | 87 | 1.62 ($\pm 1.21$) | -21.77 ($\pm 11.01$) |
| | H3K9me3 | Ovary | 86 | 2.42 ($\pm 2.44$) | -26.67 ($\pm 11.54$) |
| | H3K27ac | Gut | 104 | 2.55 ($\pm 2.92$) | -25.34 ($\pm 12.73$) |
| | H3K27ac | Head | 73 | 2.13 ($\pm 2.57$) | -20.63 ($\pm 11.84$) |
| | H3K27ac | Ovary | 68 | 3.66 ($\pm 4.44$) | -25.66 ($\pm 14.13$) |
| | Bivalent | Gut | 111 | 7.66 ($\pm 6.97$) | -71.04 ($\pm 33.03$) |
| | Bivalent | Head | 111 | 7.95 ($\pm 6.97$) | -73.27 ($\pm 32.27$) |
| | Bivalent | Ovary | 133 | 7.81 ($\pm 6.85$) | -63.87 ($\pm 33.26$) |

We found that TEs deplete H3K9me3 and H3K27ac in similar proportions (χ² test, *p*-value = 0.17; **Fig. 2D**, **Table 1** and **Table S2B-C**) and that histone mark depletion affects similarly across body parts (χ² tests, *p*-values > 0.1, **Table 1**, **Table S2E**). Notably, the bivalent depletion is the one with the highest effects and longest spread: 9.13kb (±7.05) and −78.1% (±31.23), respectively (Wilcoxon’s test, *p*-values < 0.001; **Fig. 2E-F** and **Table S2D**).

Finally, we found that TEs inducing the enrichment and depletion of epigenetic marks are highly body part-specific: 757 out of the 1,197 TEs that enrich and 376 out of 552 TEs that deplete exert their epigenetic effect in only one body part (**Fig. 2A and Table S2B**). Focusing on TEs that induce histone mark enrichment, the most body part-specific effects were the enrichment of H3K27ac and bivalent marks (308/345 and 285/304, respectively). In the case of TEs inducing depletion, the most body part-specific TEs are those depleting for H3K27ac and H3K9me3 (177/200 and 201/223, respectively) (**Fig. 2A and Table S2B**).

Overall, the majority of individual TE insertions induced the enrichment of H3K9me3 in the ovary and the enrichment of H3K27ac in the gut, although we also found enrichment of these histone marks in other body parts. Moreover, the epigenetic effects of TEs are not restricted to the enrichment of histone marks, as we also identified 552 TEs that deplete histone marks.

### 3. TEs enriching for the repressive mark are less dynamic than the ones enriching for the active or a bivalent state

When comparing the TE epigenetic state across human tissues and multiple stages of development, a small fraction of TEs was found to be consistently annotated with the same epigenetic state, with most of these metastable TEs being repressed, while TEs in active state were more dynamic (29). To test this in *D. melanogaster*, we focused on strain-specific TEs that induce the enrichment of a histone mark in more than one body part (429 TEs), and quantified how many of these TEs maintained a consistent effect, *i.e.*, were metastable, and how many of them displayed dynamic changes between body parts. We found 178 metastable TEs, from which 84% (150/178) induced the enrichment of the H3K9me3 repressive mark across all body parts (**Fig. 3**). On the other hand, among the 251 dynamic TEs, the most frequent switch is between H3K27ac and H3K9me3 (100 TEs) and between a bivalent state and H3K9me3 (95 TEs). Our results are thus in agreement with the previous results found in humans (29).

**Figure 3.**
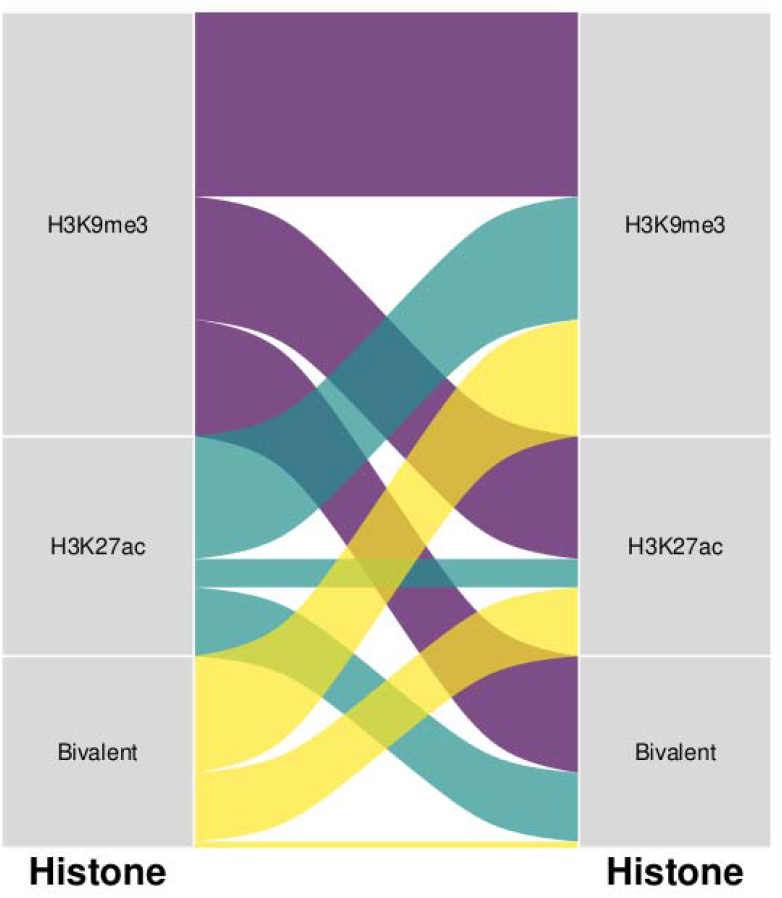
Quantification of metastable and dynamic TEs. For 429 strain-specific TEs, it was quantified which of them exhibited consistency across different body parts (metastable), and which ones underwent changes (dynamic).

### 4. LTR families are enriched for H3K9me3 while TIRs and LINE families are enriched for H3K27ac

Next, we wanted to test whether there is variation in the epigenetic effects depending on the TE family. Focusing on the enrichment of H3K9me3, we found that 58% of the TEs were LTRs (476/816), similar to the proportion previously found in *Drosophila* (15), followed by LINEs (25%, 204/816) and TIRs (17%, 136/816) (**Fig. S3A**). In the case of the enrichment of H3K27ac, we observed that overall LTRs TEs were the most common (43%, 149/345), followed by LINEs (34%, 119/345) and TIRs (23%, 81/345) **(Fig. S3B**). In the case of bivalent enrichment, the proportion is similar to that of H3K9me3 enrichment: 56% are LTRs (170/304), 25% LINEs and 19% TIRs (**Fig. S3C**).

At the family level, and focusing on those families with >20 copies, we found that TE families associated with the enrichment of H3K9me3 and those associated with the enrichment of the two marks were all LTRs: *blood*, *mdg3, Blastopia, Burdock, Copia* and *412* (**Fig. 4A-C**). On the other hand, a LINE (*Doc*) and a TIR (*H*) were the only families enriched for H3K27ac (**Fig. 4B**). Note that there are more copies than expected of TEs of the *H* family associated with H3K27ac enrichment but less with H3K9me3 enrichment (**Table S3**). Overall, LTR families are enriched for H3K9me3 and for both histones marks, while TIRs and LINEs families are enriched for H3K27ac.

**Figure 4.**
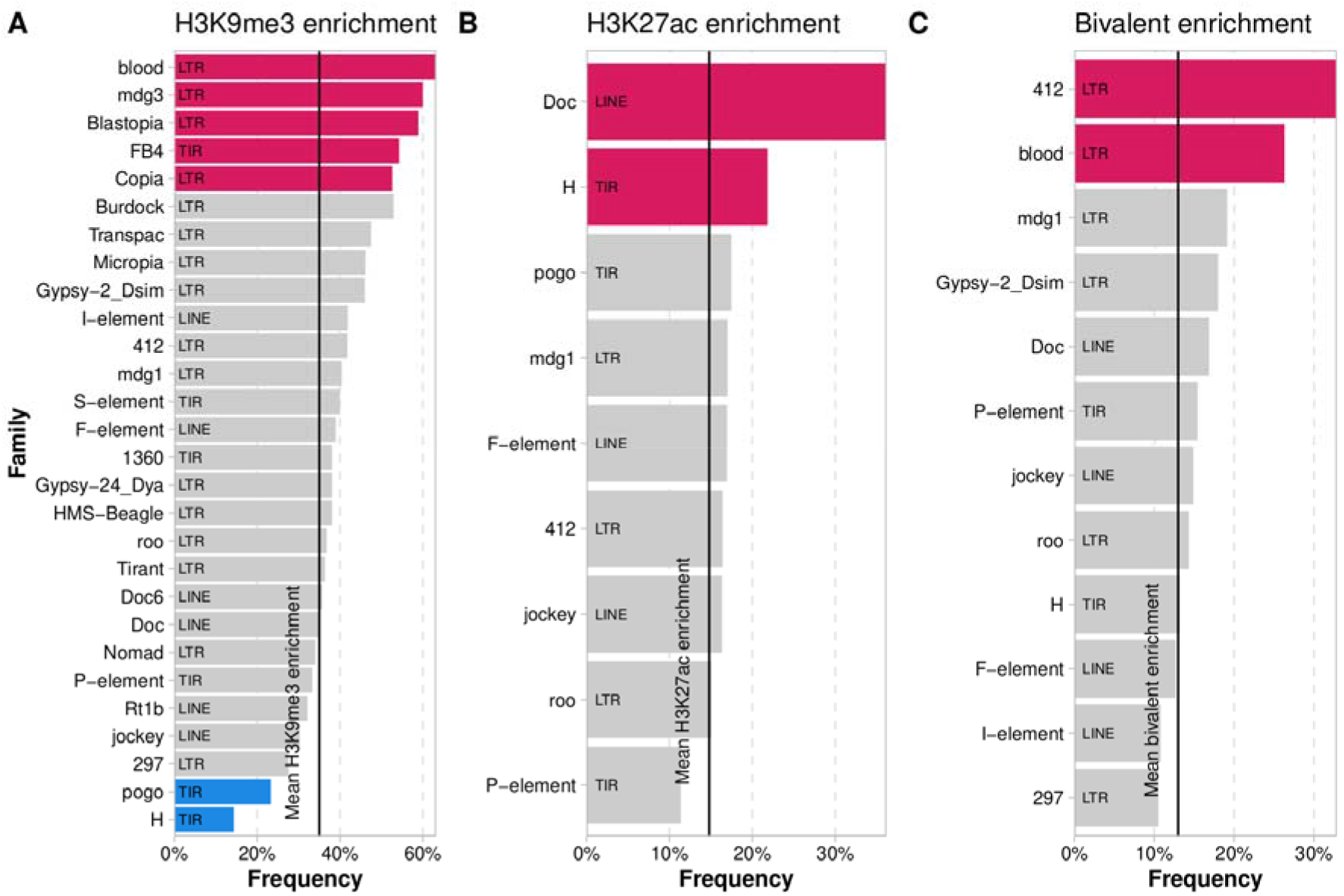
Enrichment and depletion analysis of TE families for TEs that induce enrichment of epigenetic marks. **A.** TE families enriched and depleted associated with H3K9me3 enrichment. **B.** TE families enriched and depleted associated with H3K27ac enrichment. **C.** TE families enriched and depleted associated with bivalent enrichment. Red: more copies associated with enrichment of histone mark than expected, blue: less copies than expected, gray: non-significant. Only considered families with a minimum of 8 copies with epigenetic effects and more than 20 copies in the genome. Numbers in Table S3.

### 5. 60% of the TEs that induce epigenetic changes have an effect on the expression of adjacent genes

To assess whether the epigenetic effect of TEs has an impact on the expression of nearby genes, we analyzed matching RNA-seq data available for all the strains and body-parts (25). Out of the 1,597 TEs that have epigenetic effects, 1,168 TEs had an expressed gene located within the spread of the histone marks enrichment. We calculated the *z*-score of the gene expression change between strains with and without the TE insertion. A negative *z*-score indicates that the expression level of the gene adjacent to the TE is lower than the gene expression of the same gene when the TE is not present (see *Material and Methods*). We found that 703 TEs (60%) have a significant impact on gene expression (728 genes): 389 TEs were associated with reduced gene expression, 231 with increased gene expression and 83 were associated with increased gene expression in one body part and reduced expression in another (**Table 2**).

**Table 2.**
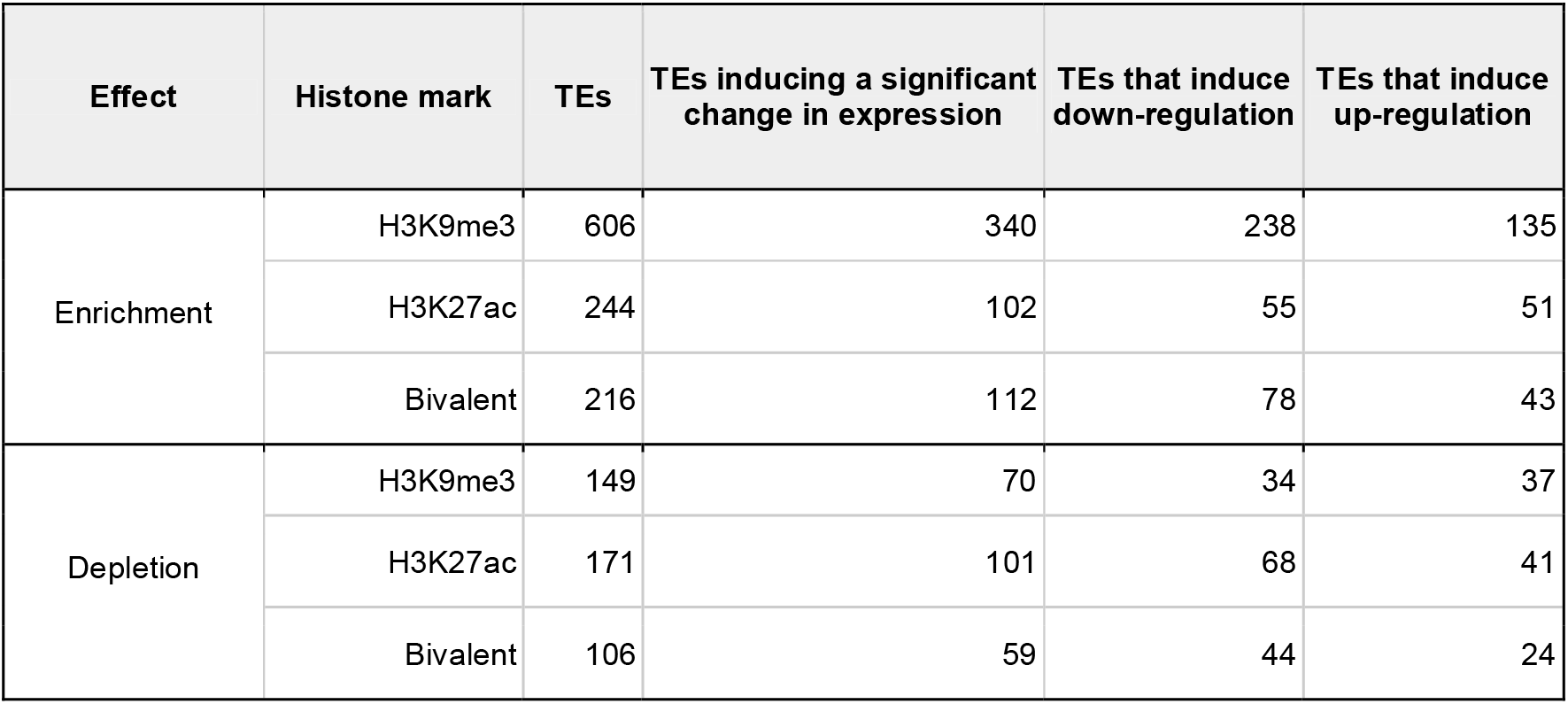
TE effects on gene expression.

We found that 70% (238/340) of the TEs that are enriched for the repressive mark are indeed associated with the down-regulation of the nearby gene (**Fig. 4A**, **Table 2** and **Table S4A**), while 50% (51/102) of the TEs that induced the enrichment of the active mark are associated with gene up-regulation (**Fig. 4B**, **Table 2** and **Table S4A**). On the other hand, the depletion of the repressive mark leads to gene up-regulation in 52% of the TEs (37/70) while the depletion of the active mark leads to gene down-regulation in 67% of the TEs (68/101). Regarding TEs that either induce bivalent enrichment or depletion, they are mostly associated with gene down regulation (70% 78/112, and 75% 44/59, respectively).

The position of the TE relative to the gene (upstream, downstream, exon or intron) was not associated with the direction of the expression change observed for H3K9me3 nor H3K27ac enrichment and depletion (**Table S4B**). The only significant association was found for the bivalent enrichment, in which TEs that induce gene down-regulation are more often located in exons (post-hoc χ² test p-value = 0.042, **Table S4B**).

Head and ovary were the body parts with more genes being significantly up or down-regulated due to the epigenetic effects of TEs compared to gut (head: 47% 228/482, gut: 41% 305/746, and ovary: 49% 427/879, χ² test *p*-value = 0.031 and 0.002, respectively).

Finally, we did not find differences on the histone mark spread, percentage of enrichment, TE position (upstream, downstream, exonic and intronic), TE length distribution, and TE frequencies between TEs with epigenetic effects inducing significant changes in expression and TEs that do not (**Table S4C**). This result suggests that the epigenetic effect of a TE, or lack thereof, is not associated with the TE itself but rather with the genomic context where it is inserted.

### 6. 23 of the TEs affecting gene expression through epigenetic changes are likely to be adaptive

TEs that are enriched for the repressive histone mark and are associated with gene down-regulation are enriched for low frequent insertions, as expected if this TEs have deleterious effects (15) (Fisher’s exact test, *p*-value < 0.001, **Table S4D**). However, nine of these TEs are present at high frequencies suggesting that they might be adaptive, with two of them previously identified as candidate adaptive TEs (Table S4E). These TEs are located nearby 11 genes related to *transport/localization* (5/11) *cell organization/biogenesis* (4/11) and *cell cycle proliferation* (3/11) (**Table S4E**). Additionally, one TE (*2R_14883304_14883308_BS2*) is associated with three cytochrome P450 genes (*Cyp6a20, Cyp6a21* and *Cyp6a9*) which have been already associated to insecticide resistance in *Drosophila* (31).

Two of the 51 TEs that are enriched for the active histone mark and associated with gene up-regulation are also present at high frequencies (**Table S4E**). The three genes associated with gene up-regulation, *Bin1*, *sra* and *Svil*, are all related to *Cell organization/biogenesis*. One TE (*FBti0019386*) affecting both *Bin1* and *sra*, was already described to be associated with gene up-regulation in cold-stress conditions and in response to infection in *D. melanogaster* (32, 33). The other TE, associated with *Svil* up-regulation has been previously identified as a candidate adaptive TE (Table S4E).

If we focus on TEs inducing depletion of histone marks, for the 37 depleting H3K9me3 that induce gene up-regulation, seven are present at high frequencies, with 5 of them involved in *Cell organization/biogenesis* (**Table S4E**). One of these, *FBti0019170*, affecting the *kuzbanian* gene, has already been associated with adaptation in response to zinc-stress tolerance (34, 35) (**Table S4D**). While *FBti0019386* was associated with H3K27ac in the gut in two of the strains analyzed, we found that this TE is also associated with H3k9me3 depletion in the gut in another strain, further suggesting that the epigenetic effect of TEs is context dependent. Additionally, two of the seven TEs were previously identified as candidate adaptive TEs (Table S4E).

Among the 68 TEs inducing the depletion of the active mark leading to gene down-regulation, there are 6 TEs present at high frequencies, associated to only two genes with a known function: one gene is a small nuclear RNA and the other one is a rho GTPase activating protein (**Table S4E**). Three of these TEs were previously identified as candidate adaptive TEs (Table S4E).

Overall, we identified that 23 of the TEs affecting gene expression through epigenetic changes are present at high population frequencies. For 10 of these 23 TEs there is previous evidence suggesting that they are evolving under positive selection, suggesting that these 23 TEs are likely candidates to be involved in adaptive processes.

### 7. Genes regulated by TEs with epigenetic effects have functions related to the neural system

To analyze whether TEs with epigenetic effects are regulating genes with similar functions, potentially indicative of a re-wiring of specific gene regulatory networks, we performed a Gene Ontology (GO) enrichment analysis (36). We found that genes up- and down-regulated in the head were involved in processes related to neuron differentiation and development (e.g.: *neuron projection* and *axonegenesis*; **Tables S5A-F** and **Fig. S4**). In the gut, there was an enrichment of neural-related terms, and also genes involved in *chemotaxis*, *cell recognition and regulation of locomotion* functions (**Tables S5D-F** and **Fig. S4**). In the ovary, although being the body part with more genes regulated by TEs, no biological processes were enriched (**Tables S5G-I** and **Fig. S4**). Finally, genes regulated by metastable TEs, *i.e.*, TEs annotated with the same epigenetic state across body parts, were also enriched in *axogenesis* and *axon development*, and also for *cell adhesion*, *syncytium formation, myoblast fusion and myotube differentiation* (**Table S5J** and **Fig. S4**).

## DISCUSSION

In this work, we have assessed the epigenetic effects of 2,235 polymorphic TEs across three *D. melanogaster* adult body parts (**Figure 1B**). The availability of high-quality TE genome annotations and ChIP-seq data for three body parts and five genomes, has allowed us to determine that 1,197 of the TEs analyzed induced the enrichment of repressive or active histone marks rather than being inserted in regions already enriched for these marks (**Figure 2**). Including RNA-seq data has also allowed us to investigate the functional consequences of TE insertions on gene expression. In contrast to previous studies (5, 15), we have found evidence for the significant influence of TEs on neighboring gene expression. We observed that 56% (340/606) of the TEs that were enriched for H3K9me3 were associated with changes in expression of genes located within the spread of this histone mark (**Figure 5 and Table 2**). Moreover, 70% (238/340) of these TEs were associated with gene down-regulation. While the previous evidence for an association between H3K9me3 enrichment and gene down-regulation in *D. melanogaster* was limited (5, 15), these studies were restricted to the analysis of TE insertions annotated in a single genome. In contrast, we have analyzed *de novo* TE annotations in five genomes, which represents a less biased dataset. Our results highlight the importance of analyzing several reference genomes to get a more comprehensive understanding of the functional consequences of TEs (26).

**Figure 5.**
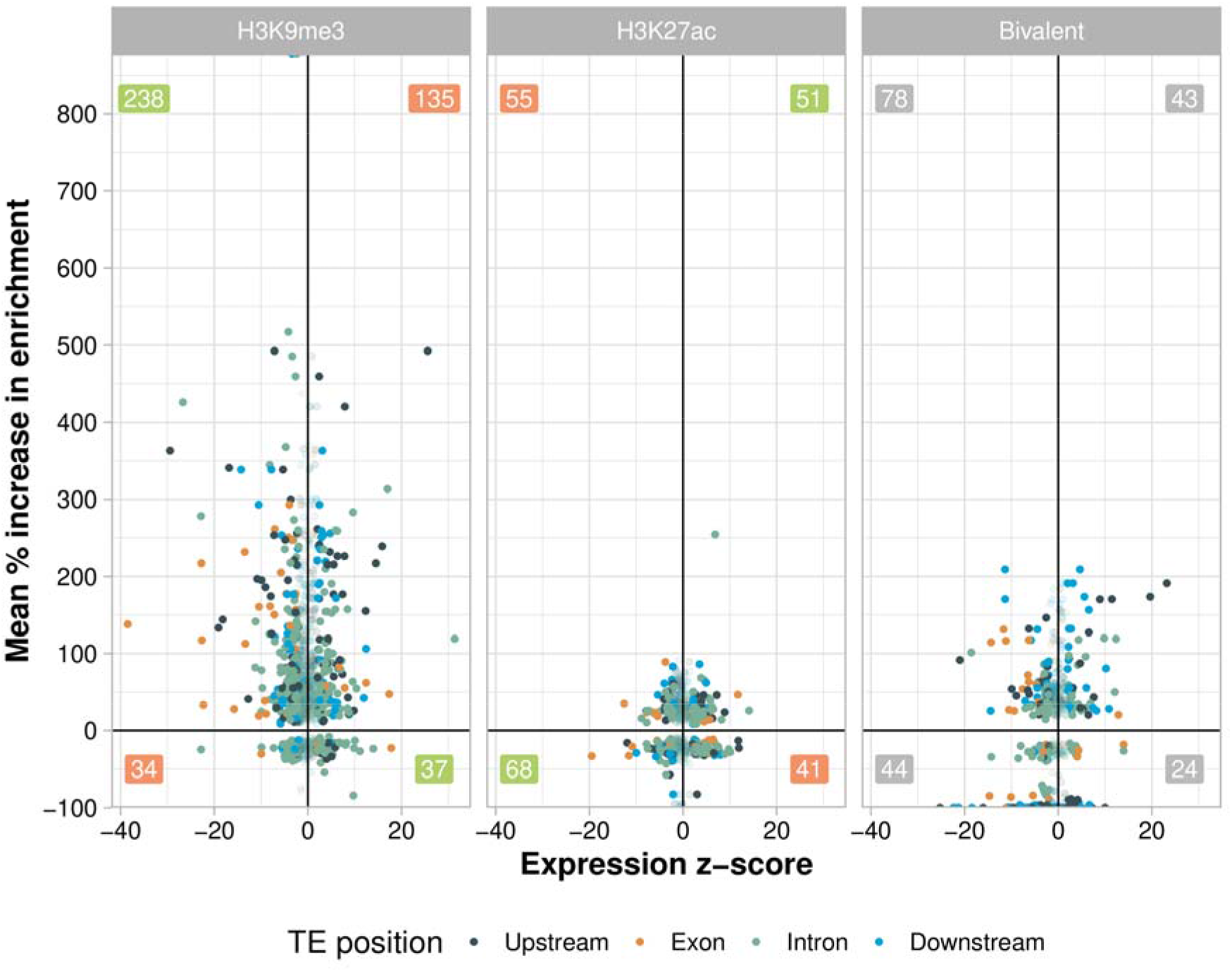
H3K9me3, H327ac and bivalent enrichment and expression *z*-score for genes with and without adjacent TE insertions. The *z*-score for gene expression of gene(TE+)-gene(TE-) (x-axis) and mean percentage of increase of the histone mark enrichment (y-axis) is plotted for pairs of TE-gene. Numbers indicate TE-gene pairs, green when the association between the epigenetic mark and the expression level is the expected, red when the association is the opposite, and gray when there are no expectations.

Although smaller, a substantial proportion of the TEs enriched for H3K27ac were also associated with changes of expression of nearby genes (40%: 102/244), with 50% of these TEs being associated with gene up-regulation (51/102). While the enrichment of H3K9me3 occurs through small-RNA mediated mechanisms (4–7), the capacity of TEs to enrich for active marks has been suggested to be associated with the recruitment by TEs of chromatin modifiers (20).

Furthermore, we also showed that the epigenetic effects of TEs extend beyond histone mark enrichment as 552 of the analyzed TEs induce the depletion of histone marks in at least one body part. The percentage of TEs disrupting H3K9me3 and H3K27ac marks and the number of TEs depleting these histone marks across body parts was similar (**Table 1**). This the pattern is not unexpected as the disruption of histone marks by TEs is due to the insertion of these sequences and not to host control mechanisms (see below). TEs disrupting epigenetic marks were also associated with gene expression changes. We found that 67% of the TEs inducing the depletion of the active mark were associated with gene down-regulation, and that 52% associated with the depletion of H3K9me3 induced up-regulation. Overall, both for TEs that enrich and for TEs that deplete histone marks, the percentage of TEs associated with the expected change of expression was high (Figure 5). While some TEs were associated with the opposite pattern of expression, it is known that the association between epigenetic marks and gene expression is complex (*e.g.* 36, 37).

By analyzing expression data on three distinct body parts -head, gut and ovary-we found that the epigenetic effects of TEs are body part-dependent. Overall, we have observed that the most prevalent epigenetic effect is the enrichment of H3K9me3 in ovaries (576 TEs). This is not unexpected, as TE insertions in the female germ line can compromise the genome integrity and the fitness of the offspring (7, 39, 40). On the contrary, the head body part exhibited fewer TEs enriching for repressive marks (140 TEs), which could be explained by the lower expression of *Su(var)3-9* gene, which encodes a key heterochromatin regulator of H3K9me3 (Fig. S1). The lower enrichment of TEs for H3K9me3 could be linked to a potentially higher rate of TE mobilization in the brain, as has been observed for *Drosophila* neurons (41) and in the human brain (41–43; but see also 44, 45). Thus, while our data suggest a lower repression of TEs in the head, more research is needed to elucidate whether this leads to an increase mobilization of TEs in the brain. We have also found that in the gut, the prevalent epigenetic effect of TEs is the enrichment of the active mark H3K27ac. TEs have previously been reported to be enriched in transcriptionally active chromatin (rich in acetylated histone H3) in the *Drosophila* midgut (30). Indeed, 5 of the 41 families enriched for H3K27ac in the gut were also found to be active by Siudeja et al. (30). During viral infections, the reverse transcriptase of certain TEs is used to generate copies of the invading virus, amplifying the immune response against viral infections (47). A higher expression of TEs in the gut, induced by H3K27ac, might thus be seen as a trade-off between TEs activity and response to viral infections occurring in the gut (47).

Overall, our study provides evidence that the epigenetic effect of TEs goes beyond gene silencing and shows that TEs can also affect gene expression by disrupting the epigenetic marks of the regions where they insert. Moreover, while genes affected by the epigenetic effect of TEs in ovary are not enriched for any biological processes, which suggests that in the ovary enrichment of TEs in repressive marks reflects the need to protect the germ line from the mutational activity of insertions, TEs in head and gut affect the expression of genes enriched for particular biological functions, suggesting that they may play a role in the rewiring of particular gene regulatory networks (Fig S4). By comparing several TE-related metrics between TEs with epigenetic effects that affect and don’t affect gene expression, we concluded that the epigenetic consequences of TE insertions are context dependent. Furthermore, 23 TEs with epigenetic effects that induce a change in gene expression were present at high population frequencies, with 10 of these TEs previously linked to an adaptive phenotypic effect or reported in genome-wide screenings for adaptive signals (Table S4E). Because the majority of TEs inducing epigenetic changes were found at low population frequencies, the analysis of more genomes, and thus of an even more complete set of TEs present in natural populations, should help elucidate the overall role of TEs in generating gene regulatory novelty by inducing or disrupting epigenetic states.

## MATERIAL AND METHODS

### Strain genomes and TE annotations

We have used five *de novo* reference genomes of five European *D. melanogaster* natural strains from the European *Drosophila* Population Genomics Consortium (DrosEU) (AKA-017, JUT-011, MUN-016, SLA-001 and TOM-007) (26). For these genomes, we have available a high-quality annotation of transposable elements (26). In brief, we only analyzed 4,823 TEs annotated in euchromatic regions and that were non-nested (**Table S6**). Genomes and TE annotations are available for visualization at the DrosOmics genome browser (25).

### TE breakpoints calls

To determine the TE insertion location (breakpoint) in genomes lacking a TE insertion, we focused on polymorphic TEs, a total of 3,878. We extracted the ±500bp flanking regions of a TE using *BEDOPS* (v2.4.39, 48) and *BEDtools* (v2.30.0, 49) and used *minimap2* (with parameters *-ax splice -G100k -B2 -t 4 -C5*) (v2.17, 50) to map such flanking regions in the genome lacking the TE. We called a TE breakpoint location when the mapping of the upstream, downstream and concatenated flanking region was unique, allowing a gap/overlap of ±50bp between the upstream and downstream regions. We could determine the insertion location for 2,425 (63%) of TEs. From them, we also removed non-nested/tandem TEs, thus analyzing 2,325 TEs. The ±20kb flanking region of each TE insertion, was divided into 20 non-overlapping 1kb windows respectively to perform ChIP-seq enrichment analyses.

### RNA-seq and ChIP-seq data for H3K9me3 and H3K27ac in three body parts

RNA-seq data and ChIP-seq data for H3K9me3 and H3K27ac in three body parts for five strains were obtained from Coronado-Zamora et al. (2023) (25), where a full description of the protocols used to generate these data can be obtained. Briefly, head, gut and ovary body parts of each strain were dissected at the same time. For each body part and strain, three replicates of 30 females aged 4-6 days were processed. RNA-seq library preparation was performed using the TruSeq Stranded mRNA sample prep kit from Illumina, and sequenced using Illumina 125bp paired-end reads. ChIP-seq libraries were performed using TruSeq ChIP library preparation kit. Sequencing was carried out in an Illumina HiSeq 2500 platform, producing 50bp single-end reads. Raw ChIP-seq data are available in the NCBI Sequence Read Archive (SRA) database (BioProject PRJNA643665), and can be retrieved and visualized through DrosOmics genome browser (25).

### ChIP-seq data processing

ChIP-seq reads were processed using *fastp* (v.0.20.1, 51) to remove adaptors and low-quality sequences. Processed reads were mapped to the corresponding *de novo* reference genome using the *readAllocate* function (*chipThres = 500*) of the *Perm-seq* R package (v0.3.0, 52), with *Bowtie* (v1.2.2, 53) as the aligner and the Csem program (v2.3, 54) in order to try to define a single location to multi-mapping reads. In all cases *Bowtie* was performed with default parameters selected by *Perm-seq*.

Then we used the ENCODE ChIP-Seq caper pipeline (v2, 55) in histone mode, using *Bowtie 2* as the aligner (53), disabling pseudo replicate generation and all related analyses *(chip.true_rep_only=TRUE*) and pooling controls (*chip.always_use_pooled_ctl=TRUE*). The MACS2 peak caller was used with default settings (Gaspar 2018). The signal value tracks produced by MACS2 were processed as follows. We used *bigWigToBedGraph* from UCSC (available at https://genome.ucsc.edu/goldenPath/help/bigWig.html) to transform bigWig files to BED files. Then, for each replicate, we used *bedmap* with the *mean* option (v2.4.39, 48) to obtain the average signal values in 10bp windows. Finally, the three replicates were averaged using *bedmap mean* to obtain a single track value for each sample that was used to estimate next the spread and enrichment of histone marks adjacent to TE insertions.

### ChIP-seq data analysis

From the 2,325 TEs with the flanking regions determined in genomes without the TE insertions [TE(-)], we computed two statistics when comparing pairs of TE(+) and TE(-) genomes (15): the *percentage increase or depletion of H3K9me3/H3K27ac enrichment* and the *extent of H3K9me3/H3K27ac spread*. In detail, we used Wilcoxon’s test (in R, v3.6.3, 56) to assess if the H3K9me3/H3K27ac enrichment or depletion in the *i*^th^ upstream and downstream windows differs significantly between pairs of TE(+) *vs.* TE(-) genomes. The most distant windows considered are 20kb from TE insertions. The *percentage increase or depletion of H3K9me3/H3K27ac enrichment* is the difference of median H3K9me3 enrichment between two genomes in the ±1kb windows immediately next to the TE insertion, divided by the enrichment level for the strain without TE. The *extent of H3K9me3/H3K27ac spread* is the farthest window in which the H3K9me3/H3K27ac enrichment or depletion is consecutively and significantly higher or lower in the genome with TE. When the farthest windows are different between the left and right sides of a TE insertion, we used the window closer to TE for the *extent of H3K9me3/H3K27ac spread* (to be conservative). To consider a TE for further analyses, we allowed only one different comparison across pairs of TE(+)-TE(-) genomes (that could be due to inconsistencies in the epigenetic change, no statistical enrichment or NA data, **Fig. S5**). We excluded 49 TEs that did not fulfill the criteria in none of the body parts. From the remaining 2,276 TEs, 1,597 have epigenetic effects in at least one body part and at least for one histone mark. 638 TEs do not have an effect (no enrichment) in none of the body parts and 41 have a “mix” effect (*i.e.*: enrich for one mark and deplete for the other) and were not analyzed. Therefore, the final dataset analyzed was of 2,235 TEs **(Fig. S6**).

### Analysis of histone enrichment patterns in TE-free regions

For every strain-specific TE insertion with epigenetic effects (2,171 TE), we selected an euchromatic region without a TE insertion annotated within 1kb. These TE-free regions were also matched to the TE analyzed regions regarding the position of the nearby gene, *i.e.,* intergenic, upstream, downstream, intron, CDS or 3’/5’UTR. Note that some of the TE insertions have more than one body part-dependent effect or affect more than one gene, we thus selected as many TE-free regions as TE associated regions affected. In total, we analyzed 5,088 null regions. We applied a Wilcoxon’s test to compare the spread and average enrichment of TE-free regions *vs.* TEs with epigenetic effects, and a χ² test to see if there is a higher proportion of TEs with epigenetic effects than null regions (in R, v3.6.3, 56) (Table S2A).

### RNA-seq data processing to obtain gene expression levels

To obtain gene expression levels from RNA-seq, we used a reference-guided transcriptome assembly obtained for each strain following Pertea et al. 2016 (57). A detailed description can be found in Coronado-Zamora et al. (2023) (25). To obtain the gene count estimations normalized for TMM (trimmed mean of the M-values), we first used the *prepDE.py* script from StringTie (58) to obtain a matrix of gene counts, with parameter *-l 125*. Next, we used the *edgeR* R package (v3.28.0, 59) to create a *DGEList* object from the table of counts using the *DGEList()* function. Next, we used *calcNormFactors()* function with the TMM method to calculate the normalization factors to scale the raw library sizes. Finally, we used the function *cpm()* to obtain the normalized counts using the TMM normalization factors.

### *Z*-score calculation

We assigned each of the TEs with epigenetic effect to the gene(s) overlapping or adjacent, taking into account the spread of the histone mark. For each pair of TE-gene, a z-score was calculated as the mean of gene expression of gene with TE (averaging the three available replicates) minus the mean expression of gene without the TE, divided by the standard error of both groups of genes. A negative z-score indicates that the gene associated with the TE has lower expression compared to the gene without a TE insertion, while a positive *z*-score indicates that the gene associated with the TE has a higher expression.

### Gene ontology (GO) enrichment analysis

To perform the GO enrichment analysis, we used the *clusterProfiler* R package (v4.2.2, 35). We used as ontology the biological processes (*ont = “BP”*), and for the significance, a *p*-value and *q*-value cutoffs of 0.05. When performing a GO enrichment in a specific body part, we used the list of genes expressed in that body part (TMM > 1) as the universe set.

### TE family enrichment

For the TE family enrichment analysis, we considered TE families with a minimum of 20 copies and 8 with epigenetic effects to perform the statistical analysis. We considered a TE family as enriched when the number of copies exhibiting epigenetic effects was significantly higher than the respective average. Specifically, we used the averages of 36.5% (816/2,235), 15.4% (345/2235), and 13,6% (304/2235) as the thresholds for H3K9me3, H3K27ac, and bivalent enrichment, respectively. To test the significance of the enrichment, we applied a χ² test (in R, v3.6.3, 56).

## DATA, MATERIALS, AND SOFTWARE AVAILABILITY

All scripts and R Markdown documents to reproduce analyses and figures have been deposited in GitHub: https://github.com/GonzalezLab/epigenetic-effects-transposons-dmelanogaster.

## ACKNOWLEDGMENTS

We thank Grace Lee for discussions and suggestions on how to perform some of the analyses. This project has received funding from the European Research Council (ERC) under the European Union’s Horizon 2020 research and innovation programme (H2020-ERC-2014-CoG-647900), from grant PID2020-115874GB-I00 funded by MCIN/AEI/10.13039/501100011033 and from grant 2021 SGR 00417 funded by Departament de Recerca i Universitats, Generalitat de Catalunya awarded to J.G.

## AUTHOR CONTRIBUTIONS

JG conceived the project; JG and MCZ designed the analyses; MCZ performed the analyses; JG and MCZ wrote and revised the manuscript.

## SUPPLEMENTARY FIGURES

**Supplementary Fig. 1.**
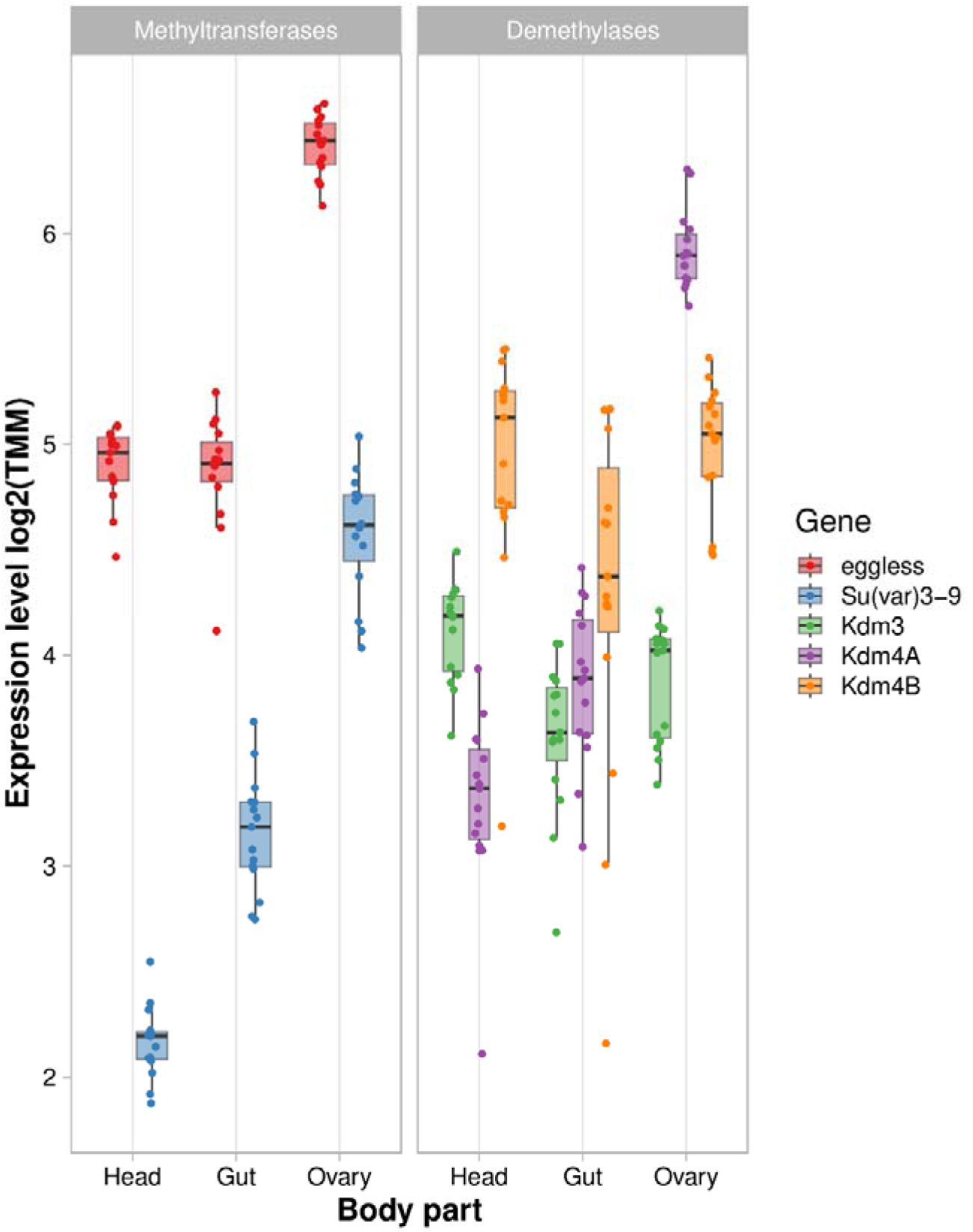
Expression level of methyltransferases and demethylases associated with H3K9me3 by body part. Expression levels in log2 (TMM). Each observation corresponds to a replicate of each strain (in total 15 observations per gene).

**Supplementary Fig. 2.**
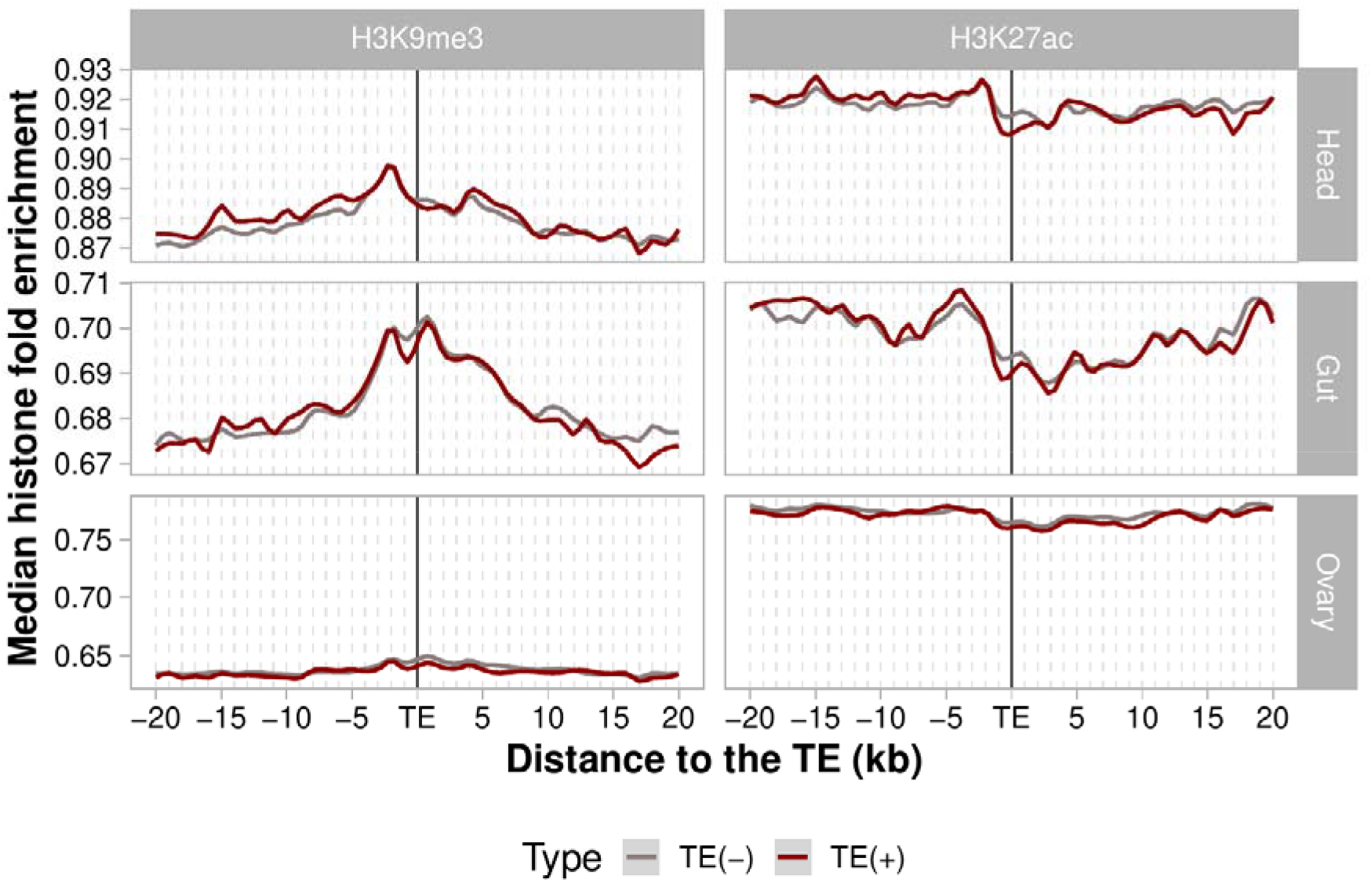
B. Median H3K9me3 and H2K27ac fold enrichment for “null” regions (*n*=5,175 regions). Red line represents histone mark enrichment of a genomes with a null region and the gray line represents histone mark enrichment of genomes without the null region. H3K9me3 and H3K27ac fold enrichment was averaged (median) over all sequences flanking the analyzed regions of all genomes. Plots were generated using LOESS smoothing (span = 10%).

**Supplementary Fig. 3.**
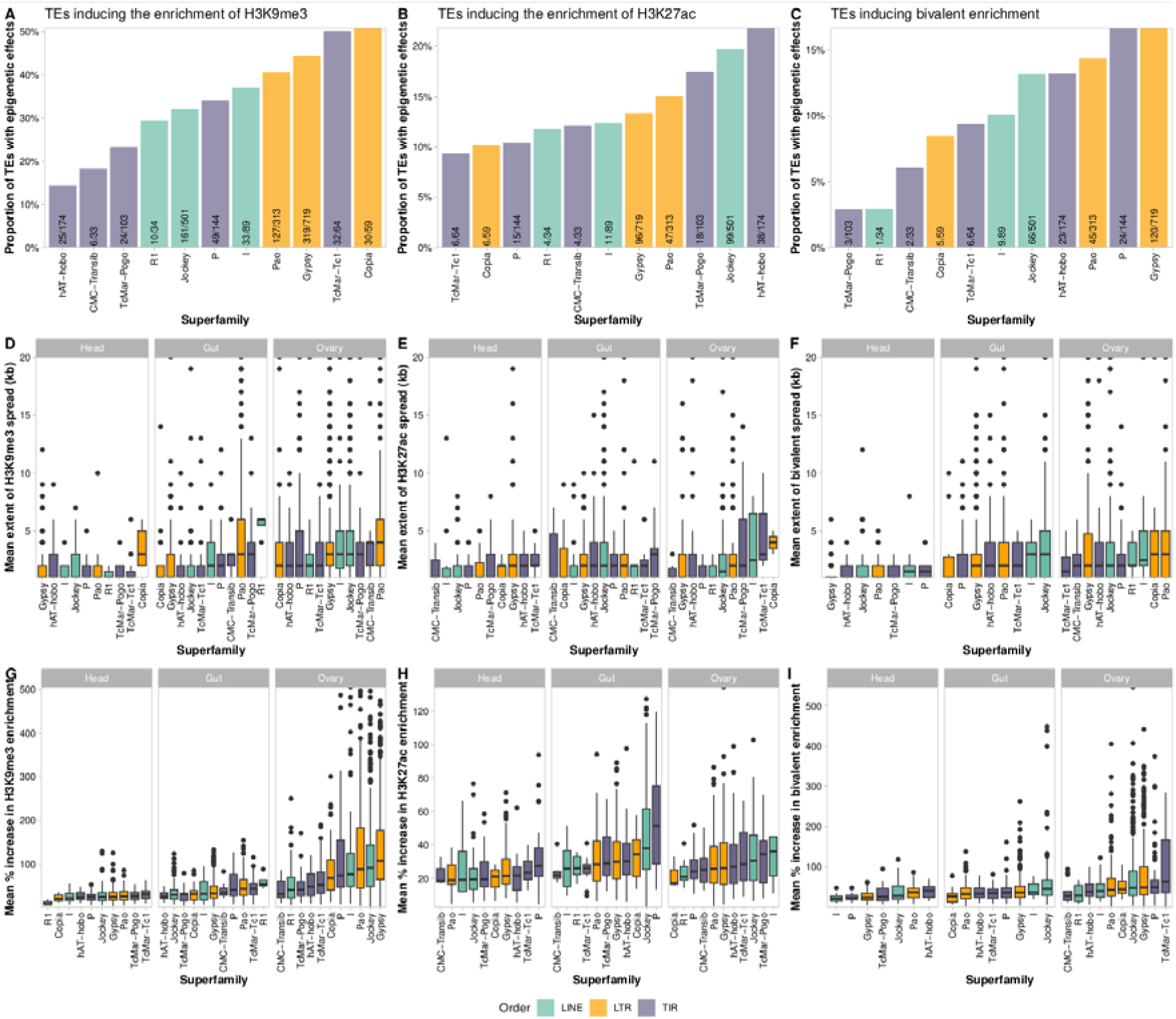
Proportion and effects of TEs that enrich for epigenetic across different TE superfamilies. **A.** TEs inducing the enrichment of H3K9me3 by superfamily. **B.** TEs inducing the enrichment of H3K9me3 by superfamily. **C.** TEs inducing bivalent enrichment by superfamily. **D.** Mean extent of H3K9me3 spread (kb) across superfamilies. **E.** Mean extent of H3K27ac spread (kb) across superfamilies. **F.** Mean extent of bivalent spread (kb) across superfamilies. **G.** Mean percentage of increase of H3K9me3 across superfamilies. **E.** Mean percentage of increase of H3K27ac across superfamilies. **F.** Mean percentage of increase of bivalent enrichment across superfamilies. Different colors denote different order of TEs (green = LINE, orange = LTR, purple = TIR).

**Supplementary Fig. 4.**
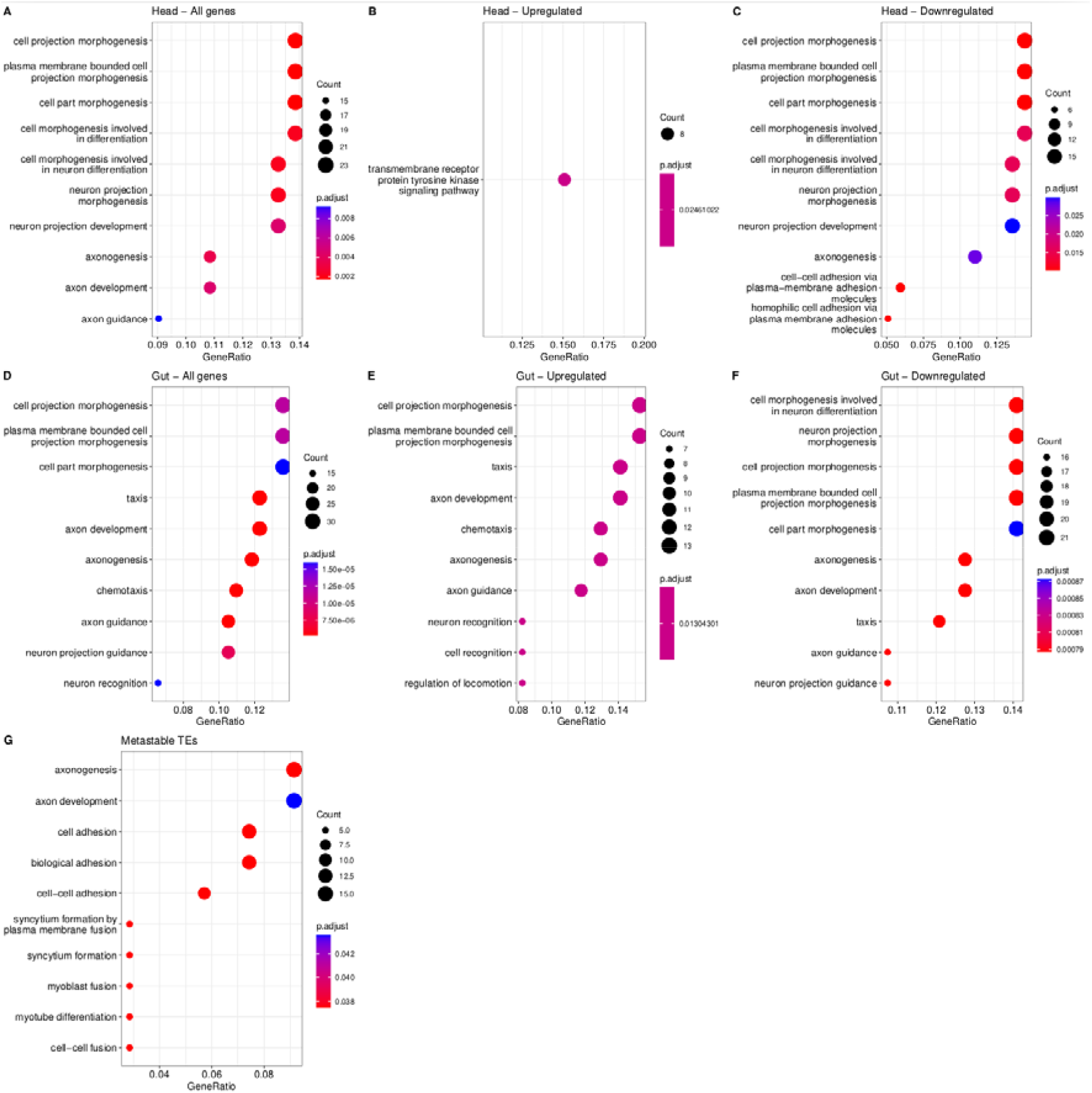
GO biological process enrichment results for genes associated to TEs with epigenetic effects. **A.** For all genes associated to TEs in head. **B.** For genes up-regulated in head. **C.** For genes down-regulated in head. **D.** For all genes associated to TEs in gut. **E.** For genes up-regulated in gut. **F.** For genes down-regulated in gut. **G.** For genes associated to metastable TEs. Data in Table S5.

**Supplementary Fig. 5.**
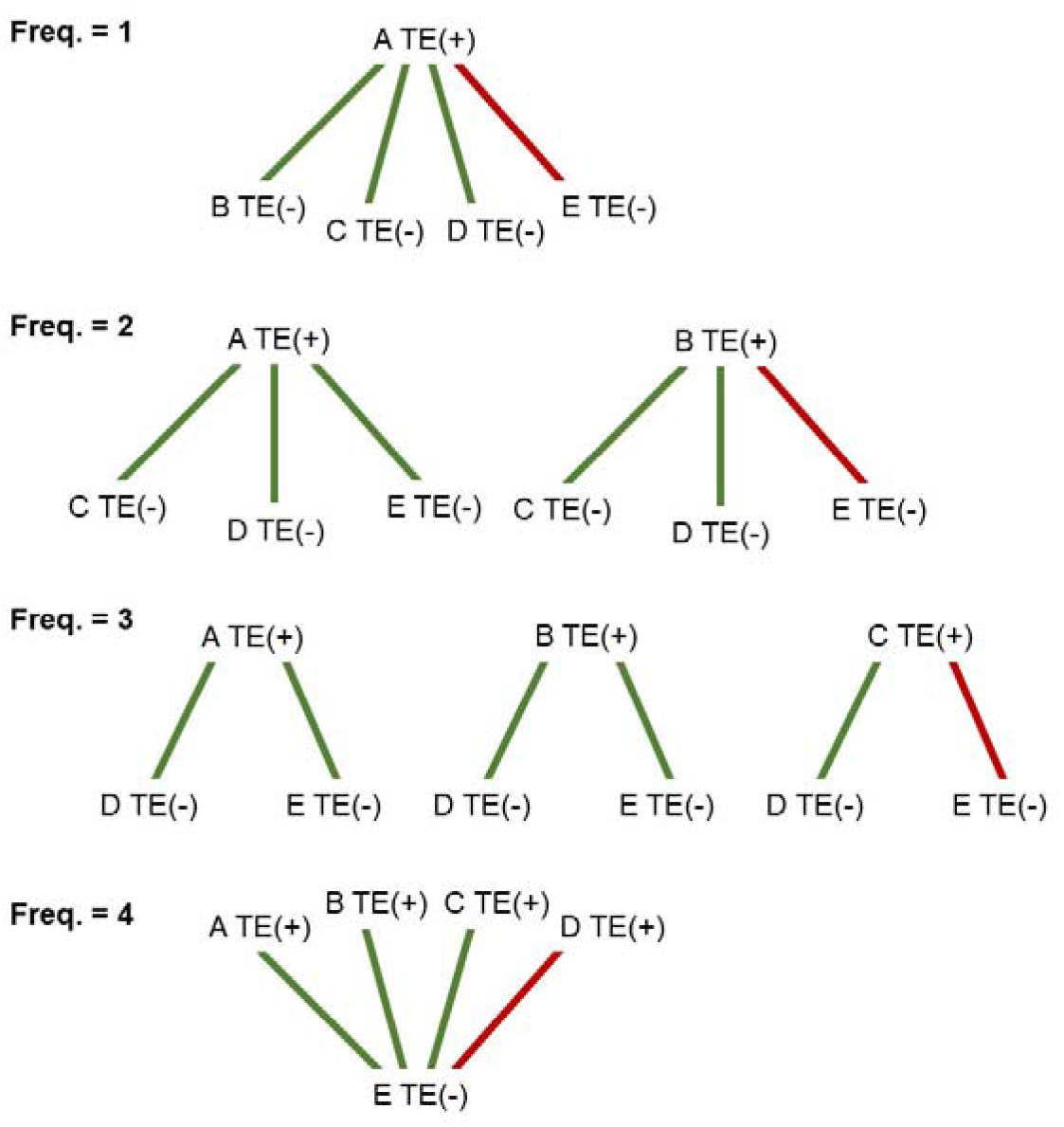
To consider a TE with epigenetic effects we allowed only one different comparison across pairs of TE(+)-TE(-) genomes. A, B, C, D and E represent strains’ names. TE(+) and TE(-) represent genomes with and without a TE insertion, respectively. Green indicates consistent comparisons, while red denotes inconsistencies compared to the most prevalent change.

**Supplementary Fig. 6.**
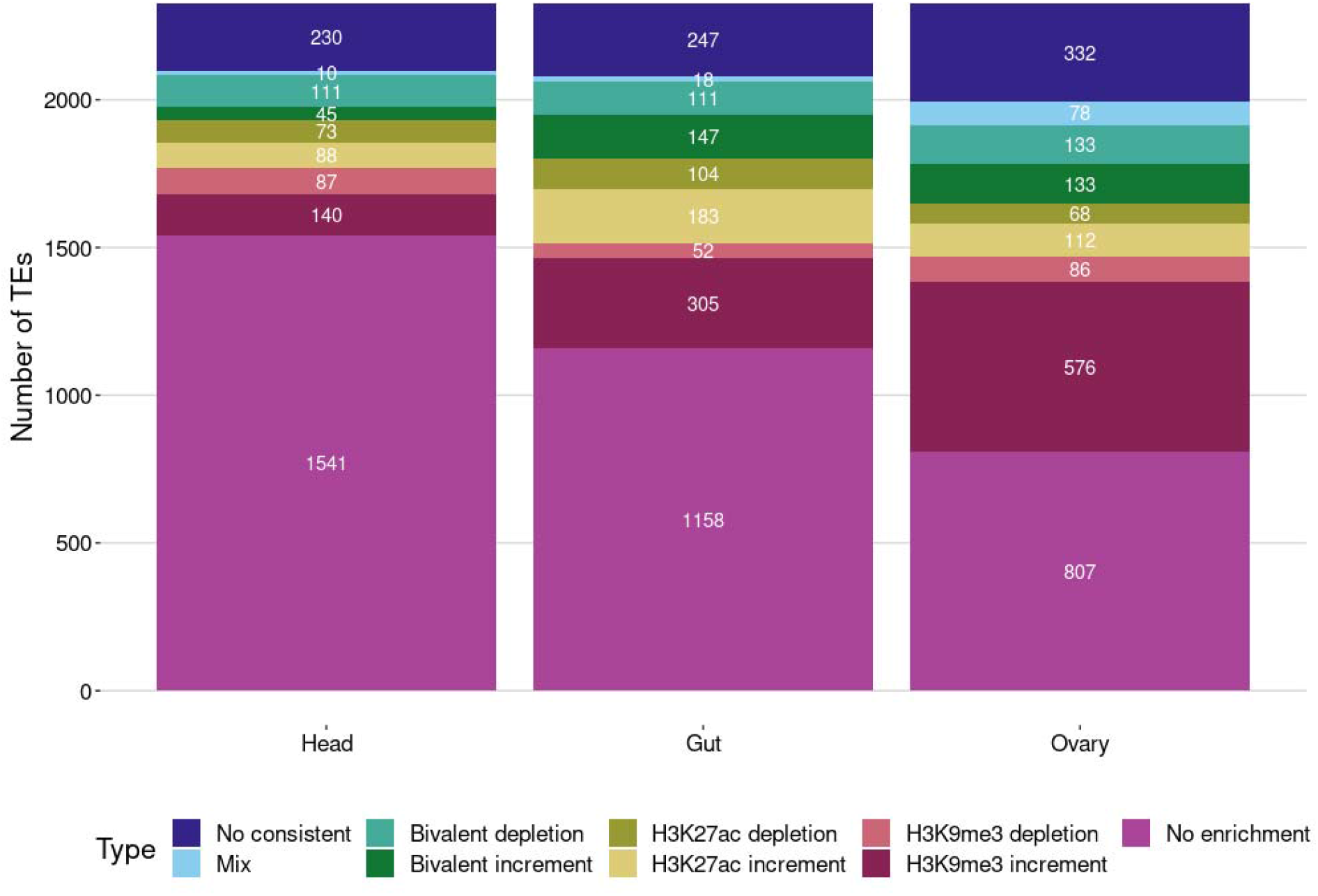
Status of 2,325 polymorphic TEs across body parts.

